# Multifocal tDCS modulates resting-state functional connectivity in older adults depending on induced electric field and baseline connectivity

**DOI:** 10.1101/2020.05.15.090860

**Authors:** Kilian Abellaneda-Pérez, Lídia Vaqué-Alcázar, Ruben Perellón-Alfonso, Cristina Solé-Padullés, Núria Bargalló, Ricardo Salvador, Giulio Ruffini, Michael A. Nitsche, Alvaro Pascual-Leone, David Bartrés-Faz

## Abstract

**Background:** Advancing age affects the brain’s resting-state functional networks. Combining non-invasive brain stimulation (NIBS) with neuroimaging is a promising approach to modulate activity across resting-state functional systems and explore their true contribution to cognitive function in aging. However, substantial individual variability in the response to NIBS has been reported and, hence, identifying the individual predictors of NIBS-induced modulatory effects is crucial if we are to harness their potential.

**Methods:** Thirty-one cognitively healthy older adults (71.68 ± 2.5 years; 19 females) underwent two different multifocal real tDCS conditions (C1 and C2) and a sham condition in a crossover design during a resting-state functional magnetic resonance imaging (rs-fMRI) acquisition. The real tDCS conditions were designed to induce two distinct electric field distribution patterns either targeting generalized cortical overactivity or a dissociation between the frontal areas and the posteromedial cortex. Stimulation was delivered through an MRI-compatible device using 8 small circular electrodes. Each individuals’ anatomical T1-weighted MRI was used to generate a finite element model to define the individual electric field generated by each tDCS condition.

**Results:** The two tDCS conditions modulated resting-state connectivity differently. C1 increased the coactivation of numerous functional couplings as compared to sham, however, a smaller amount of connections increased in C1 as compared to C2, while no differences between C2 and sham were appreciated. At the group level, C1-induced modulations primarily included temporo-occipital areas and distinct cerebellar regions. This functional pattern was anatomically consistent with the estimated distribution of the induced electric field in the C1 condition. Finally, at the individual level, the extent of tDCS-induced rs-fMRI modulation in C1 was predicted by baseline resting-state connectivity and simulation-based electric field magnitude.

**Discussion:** Our results highlighted that multifocal tDCS procedures can effectively change neural dynamics in the elderly consistently with the spatial distribution of the estimated electric fields on the brain. Furthermore, we showed that specific brain factors that have been revealed to explain part of the individual variability to NIBS in young samples are also relevant in older adults. In accordance, designing multifocal tDCS configurations based on specific fMRI patterns appears to be a valuable approach to precisely adjust those complex neural dynamics sustaining cognition that are affected as a function of age. Furthermore, these innovative NIBS-based interventions should be individually-tailored based on subject-specific structural and functional data to ultimately boost their potential in aged populations.

## 1. Introduction

The human brain is organized into complex neural networks that can be studied through resting-state functional magnetic resonance imaging (rs-fMRI; Damoiseaux et al., 2006; Petersen & Sporns, 2015; Smith et al., 2009; van den Heuvel & Sporns, 2013). Some of these neural systems have been shown to support cognition (Bressler & Menon, 2010) and change through the lifespan (Betzel et al., 2014; Dosenbach et al., 2011), as well as to be highly susceptible to aging (Ferreira & Busatto, 2013; Nashiro et al., 2017; Sala-Llonch et al., 2015; Tomasi & Volkow, 2012).

Non-invasive brain stimulation (NIBS) protocols have been used to modulate these cognitive systems in older adults (Abellaneda-Pérez et al., 2019a). The combination of NIBS with neuroimaging techniques in these populations has shed light into the network plasticity mechanisms underlying cognitive maintenance and reserve in the elderly (i.e., Abellaneda-Pérez et al., 2019b) as well as on the putative neurobiological mechanisms underlying NIBS-induced phenotypic improvements in aged samples (i.e., Antonenko et al., 2018; Holland et al., 2011; Meinzer et al., 2013; Nilakantan et al., 2019).

Transcranial direct current stimulation (tDCS) has been widely employed in the latest years, particularly in advancing age (i.e., Perceval et al., 2016; Tatti et al., 2016). In conventional tDCS studies, which aim to target discrete cortical regions, a single anode accompanied by its corresponding cathode is used. According to our current mechanistic understanding, during tDCS, neural membrane potentials are depolarized under the anode, leading to an increase in cortical excitability, while they are hyperpolarized under the cathode, thus diminishing cortical excitability at the macroscopic level (Nitsche & Paulus, 2000; Nitsche et al., 2008). More recently, novel multifocal or network-based tDCS protocols have been developed in order to target multiple brain areas simultaneously (Ruffini et al., 2014). In multifocal tDCS, multiple electrodes with differential intensities and polarities are employed such that the resulting field aims to maximally target a specific distributed brain network and can result in higher modulatory efficacy than an otherwise similar two-electrode tDCS approach in younger samples (i.e., Fischer et al., 2017). However, the effects of multifocal tDCS on aged populations remains unexplored, despite its potential to modulate a complex system rather than a concrete region. In this vein, a whole network modulation is particularly relevant in aging, since, as observed in numerous descriptive fMRI investigations, brain networks are less integrated and more segregated as a function of age (Cao et al., 2014; Chan et al., 2014; Grady et al., 2016; Sala-Llonch et al., 2014; Spreng et al., 2016), perhaps reflecting age-associated dedifferentiation processes (Park et al., 2012). Notably, this loss of brain functional segregation has been related with poorer cognitive performance in aging (i.e., Sala-Llonch et al., 2014). Consequently, altering multiple neural nodes within or between specific neural circuits could be used to restore a regular brain functioning and thus ameliorate cognition in advanced age.

The present study draws from a previous investigation by our group and is based on its fMRI findings. In the stated report, we observed that older adults can engage distinct brain activity patterns to successfully solve a particular working memory paradigm (see Fernández-Cabello et al., 2016 for further details). In the present study, two real tDCS conditions were designed based on those previous fMRI findings, to induce two distinct electric field distribution patterns, which were expected to produce either generalized cortical overactivity (i.e., condition 1, or C1) or an antero-posterior dissociation aiming to enhance frontal areas whereas reducing the posteromedial cortex activity (i.e., condition 2, or C2; see section 2.4 for detailed information). The goals of the present study were: (1) to explore the impact of two distinct multifocal tDCS montages on rs-fMRI associated connectivity changes in older adults; (2) to examine the correspondence between the tDCS-induced rs-fMRI effects and the estimated electric field on the brain at the group level; and (3) to determine individual predictors of the observed rs-fMRI effects based on the individual’s baseline resting-state functional connectivity (rs-FC) and the estimated electric field magnitude.

## 2. Materials and methods

### 2.1 Participants

As in our previous studies (Vaqué-Alcázar et al., 2017; 2020; Vidal-Piñeiro et al., 2014), subjects participating in the present investigation were recruited from the Fundació Institut Català de l’Envelliment. We contacted thirty-seven subjects and initially included thirty-three participants fulfilling the following criteria: aged older than 65 and neuropsychological assessment within range of normality (see below). Selected exclusion criteria were Hamilton Depression Rating Scale >13, history of epilepsy, neurological or psychiatric disorders and any NIBS-related contraindication (Antal et al., 2017; Rossi et al., 2009) as well as for the MRI. One subject was discarded for a previous single episode of absence seizure and another volunteer was excluded due to morphine pump implantation to treat chronic pain.

Finally, thirty-one subjects met criteria to participate in this study. All participants were tDCS naïve and right-handed older adults [mean age ± standard deviation (*SD*), 71.68 ± 2.5 years; age range, 68 – 77 years; 19 females; years of education mean ± *SD*, 12.29 ± 4.0 years]. All volunteers provided informed consent in accordance with the Declaration of Helsinki (1964, last revision 2013). All study procedures were approved by the Institutional Review Board (IRB 00003099) at the University of Barcelona. For all participants, MRI images were examined by a senior neuroradiologist (NB) for any clinically significant pathology (none found).

### 2.2 Experimental design

The present study was conducted in a randomized sham-controlled crossover design that consisted of four visits to our center. On the first visit (i.e., pre-experimental session; day 0), all participants underwent a comprehensive neuropsychological assessment to ensure cognitive functioning within the normal range according to age and years of education (see section 2.3). Subsequently, three experimental sessions with distinct multifocal tDCS conditions (days 1, 2 and 3) were conducted while acquiring rs-fMRI during brain stimulation. In addition, on day 1, a high-resolution three-dimensional (hr-3D) dataset was acquired for functional data preprocessing purposes and to estimate the distribution and magnitude of the induced electric field. In the experimental sessions, multifocal tDCS was applied to all participants using the two different montages refered above (i.e. C1 and C2) with real stimulation as well as a sham stimulation condition (see section 2.4 for further details). The order of the experimental sessions was counterbalanced. There was a minimum one-month wash-out between each stimulation session to avoid possible prevalence of after-effects (Fig. 1).

**Fig. 1.**
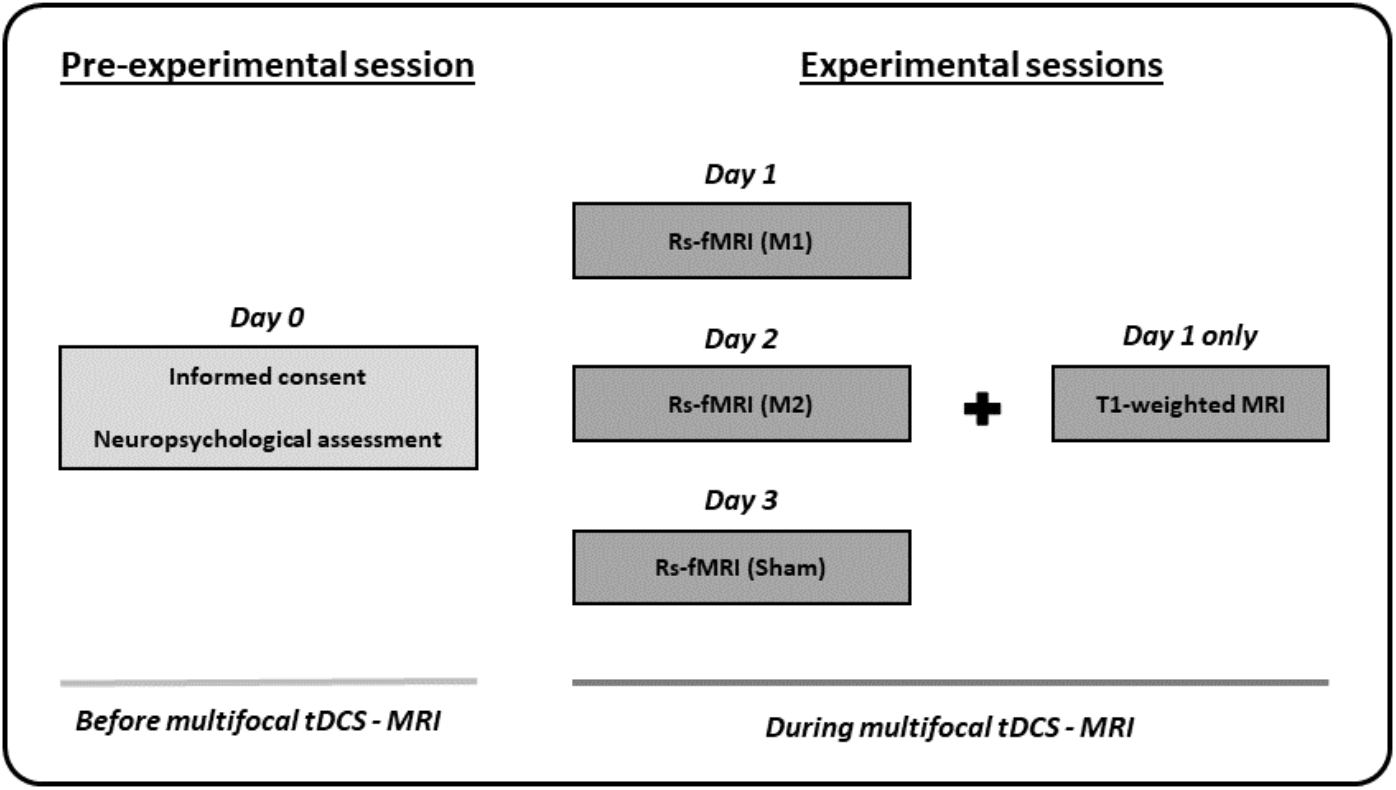
Study protocol and timeline of the procedures accomplished before and during multifocal tDCS-MRI. Abbreviations: tDCS, transcranial direct current stimulation; MRI, magnetic resonance imaging; rs-fMRI, resting-state functional MRI.

### 2.3 Neuropsychological assessment

A comprehensive battery of neuropsychological tests covering all cognitive domains was administered (see Supplementary Material [SM] for detailed information). All participants presented a normal cognitive profile with mini-mental state examination (MMSE) scores of ≥ 27 and performance scores not more than 1.5 *SD* below normative data (adjusted for age and years of education) on any of the administered neuropsychological tests (i.e., they did not fulfill the criteria for mild cognitive impairment; Petersen & Morris, 2005).

### 2.4 tDCS parameters

Two distinct multifocal tDCS montages were designed with the Stimweaver montage optimization algorithm (Ruffini et al., 2014). The latter determines the positions and currents of the electrodes over the scalp that induce an electric field in the brain that better approximates a weighted target electric field map. In this optimization we optimized for the electric field component normal (orthogonal, *E*_*n*_) to the cortical surface, assuming a first order model for the interaction of the electric field with neurons in the cortex: when *E*_*n*_ points into/out of the cortical surface (positive/negative values of *E*_*n*_ in our convention), this leads to an increase/decrease in the membrane potential of the soma of pyramidal cells (and hence, cortical excitability). As mentioned, the weighted target *E*_*n*_-maps used in this study were designed based on the findings obtained in our previous fMRI study (Fernández-Cabello et al. 2016). More precisely, on the one hand, the C1 montage was grounded on an fMRI pattern of extended cortical activity, including the bilateral middle frontal gyri, the paracingulate gyri, the precuneus cortex, the bilateral supramarginal gyri and/or intraparietal sulcus area and the lingual gyri (Fig. 2A). On the other hand, the C2 montage was derived from a second fMRI pattern including moderate activity increases in the bilateral middle frontal and the paracingulate gyri, altogether with brain activity decreases of the posterior cingulate gyrus, the ventral precuneus, and the precentral gyri (Fig. 2B). In the optimization, the regions that registered a brain activity increase/decrease were targeted with an excitatory/inhibitory (positive/negative) target *E*_*n*_ field (see Fig. S1 and S2). Cortical maps of the magnitude of the induced electric field, in which our analyses were focused, are shown in Fig. 2C for C1 and in Fig. 2D for C2, in the template head model used in the optimization (Colin27; Miranda et al, 2013). Stimulation was delivered via an MRI-compatible Starstim Neuroelectrics^®^ device, using 8 circular MRI Sponstim electrodes with an area of 8 cm^2^. The electrodes were located on the participant’s scalp by fitting them inside a sponge and into the holes of a neoprene cap corresponding to the 10/10 international system for electrode placement. The central Cz position was aligned to the vertex of the head in every subject to ensure an accurate cap placement.

**Fig. 2.**
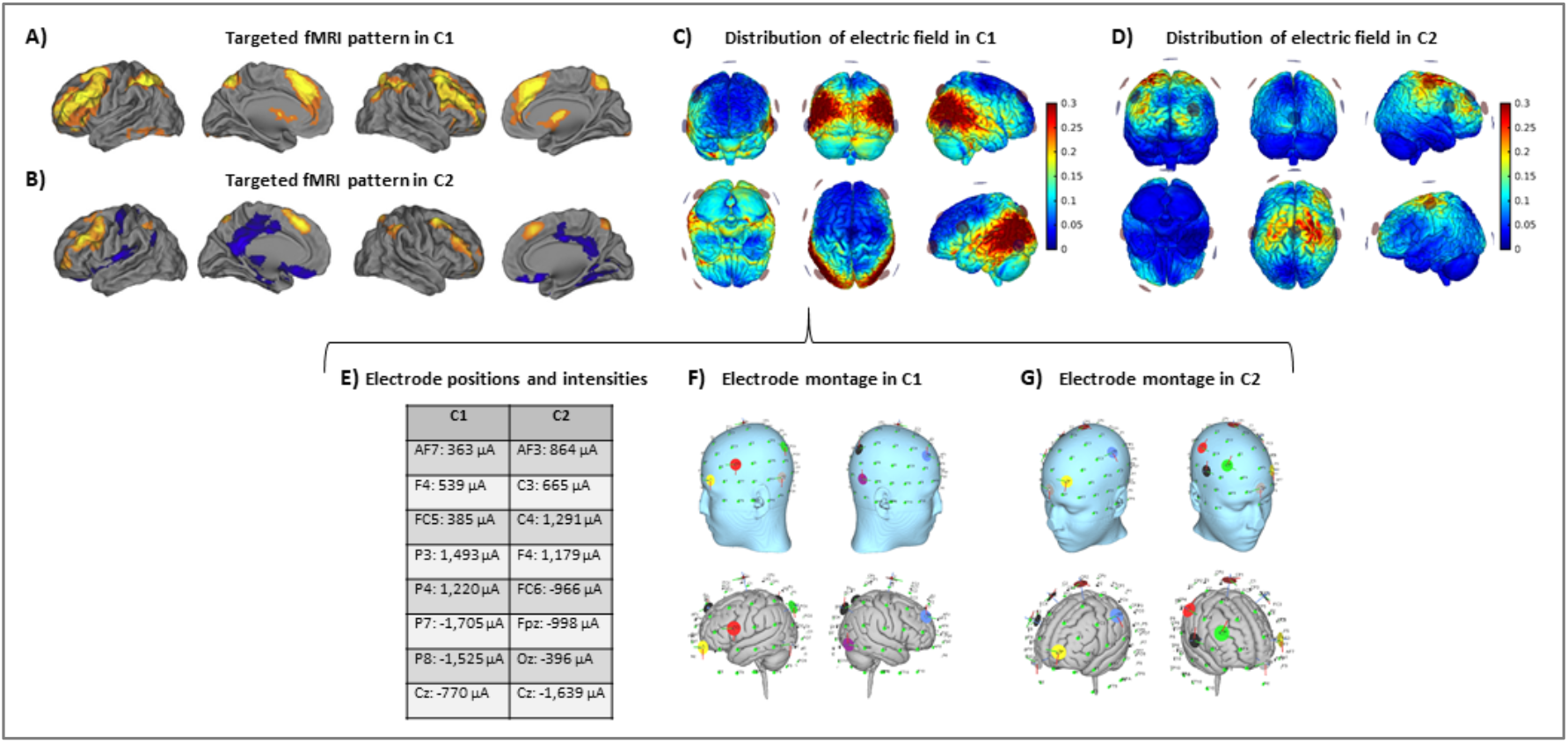
Multifocal tDCS montages. Targeted fMRI patterns in A) C1 and B) C2, from Fernández-Cabello et al. (2016, adapted with permission). Distribution of the electric field’s magnitude in the cortical surface in C) C1 and D) C2 (in V/m units). E) Electrode positions and current intensities in C1 and C2. Representation of the electrode locations and shape on the scalp (top) and the gray matter (bottom) based on a Montreal Neurological Institute (MNI) head in F) C1 and G) C2, obtained for visual purposes.

Based upon the foregoing, in the C1 montage the electrodes were placed in the AF7, F4, FC5, P3, P4, P7, P8 and Cz positions (Fig. 2E and 2F), while in the C2 montage the electrodes were placed in the AF3, C3, C4, F4, FC6, FPZ, OZ and Cz positions (Fig. 2E and 2G). For sham stimulation, either C1 or C2 montages were randomly used. The stimulator was situated outside the MRI room and the electrodes were soaked with saline solution and a thin layer of Ten20 conductive paste to ensure good conductivity and stability throughout the MRI acquisition. The current was delivered to each electrode with a wireless neurostimulator (Starstim, by Neuroelectrics Barcelona) connected to a computer via Bluetooth. For safety issues, the maximum current delivered by any electrode was 2 mA, while the maximum current injected through all the electrodes was 4 mA. In the real intervention conditions, the current was supplied during the whole rs-fMRI acquisition. In all groups, the current was initially increased and finally decreased in a 30 s ramp-up and ramp-down fashion. For the sham condition, the current dosage was composed of an initial ramp-up of 30 s immediately followed by a 1 min ramp-down, and a final ramp-down of 30 s immediately preceded by a ramp-up of 1 min.

### 2.5 MRI acquisition

All participants were scanned with a Siemens Magnetom Trio Tim Syngo 3 Tesla system at the MRI Core Facility (IDIBAPS) of the Hospital Clínic de Barcelona, Barcelona, Spain. Three identical rs-fMRI datasets (T2*-weighted GE-EPI sequence; interleaved acquisition; repetition time [TR] = 2,700 ms; echo time [TE] = 30 ms; 40 slices per volume; slice thickness = 3.0 mm; interslice gap = 15%; voxel size = 3.0 × 3.0 × 3.0 mm; field of view [FOV] = 216 mm; 178 volumes) and a hr-3D structural dataset (T1-weighted magnetization-prepared rapid gradient-echo [T1-weighted MPRAGE]; sagittal plane acquisition; TR = 2,300 ms; TE = 2.98 ms, inversion time [IT] = 900 ms; slice thickness = 1.0 mm; voxel size = 1.0 × 1.0 × 1.0 mm; FOV = 256 mm; 240 slices) were acquired.

### 2.6 Functional connectivity analyses

The FMRIB Software Library (FSL; version 6.00; http://fsl.fmrib.ox.ac.uk/fsl/fslwiki/) and the Analysis of Functional NeuroImages (AFNI; https://afni.nimh.nih.gov/) were used for preprocessing and analyzing functional neuroimaging data.

#### 2.6.1 Functional connectivity preprocessing

Rs-fMRI data preprocessing included the removal of the first five volumes, motion correction, skull stripping, spatial smoothing (Full Width at Half Maximum [FWHM] = 7 mm), grand mean scaling and filtering with both high-pass and low-pass filters (0.01- and 0.1-Hz thresholds, respectively). Data were then regressed with six rigid-body realignment motion parameters, mean white matter, and mean cerebrospinal fluid signal. No global signal regression was used. Normalization to MNI standard space was also applied. Moreover, as head movement may affect rs-fMRI results (Power et al., 2012, 2015; van Dijk et al., 2012), in-scanner head motion was calculated for every subject. More precisely, two standard measures to estimate in-scanner head motion were obtained in a similar manner as described elsewhere (Power et al., 2012). Displacement relative to a single reference volume (absolute displacement) and relative to the precedent volume (relative displacement) were calculated for every subject. In our sample, no significant differences were found between the three conditions (i.e., C1, C2 and sham), considering both absolute (*χ*^*2*^ = 1.806, *p* = 0.405) and relative displacement (*χ*^*2*^ = 2.323, *p* = 0.313).

#### 2.6.2 ROI-based functional connectivity analyses

Functional connectivity analyses were implemented based on a whole-brain atlas that parcels the brain into a set of anatomical regions of interest (ROIs). The selected atlas was the one developed for the CONN toolbox (www.nitrc.org/projects/conn, RRID: SCR_009550; Whitfield-Gabrieli & Nieto-Castanon, 2012). This atlas includes a rich set of regions to perform comprehensive whole-brain analyses using ROI-based approaches. More specifically, this atlas includes 132 ROIs, combining the FSL Harvard-Oxford cortical (91 ROIs) and subcortical atlases (15 ROIs) and the cerebellar areas from the Anatomical Automatic Labeling (AAL) atlas (26 ROIs). Individualized time-series of the different ROIs were extracted from the preprocessed and regressed images. In order to obtain a rs-FC measure for each ROI-to-ROI connection in each subject, the acquired ROI time-series were correlated with one another to create correlation matrices, using Pearson product-moment correlations.

### 2.7 Simulation of electric fields

SimNIBS 3.0.7 was used to calculate the electric fields induced by tDCS based on the FEM and individualized head models derived from the structural MRI datasets (www.simnibs.org; (Thielscher et al., 2015; Windhoff et al., 2013). First, T1-weighted anatomical images were used to create individualized tetrahedral FE head meshes of each subject, using MATLAB toolboxes (Nielsen et al., 2018), MeshFix (Attene, 2010), and Gmsh (Geuzaine & Remacle, 2009). These head models contain representations of the scalp, skull, cerebrospinal fluid (including the ventricles), eyeballs, grey-matter and white-matter. Second, the electrode positions (i.e., the center coordinates of the modeled electrodes) were placed on each subject head mesh, according to the locations established for each montage (i.e., C1 and C2). Then, electric field simulations were computed for each condition separately. The electrode shape was set as elliptical, and the size was defined as 2.3 cm of diameter and 1 mm of thickness. The electrode’s sponge size was defined as 3.2 cm of diameter and 3 mm of thickness. Tissue and electrode conductivity values were set as default in SimNIBS software (Saturnino et al., 2015; Thielscher et al., 2011). Third, individual results were averaged together for each condition, resulting in group averages and *SD*s of the electric field magnitude. Finally, for each montage, the mean of the top percentiles (99.9%) of the field and the electric field magnitude within three ROIs based on fMRI findings was computed. Specifically, these ROIs were the left and right temporo-occipital medial temporal gyri (toMTG.L and toMTG.R, respectively) and the right planum temporale (PT.R; see section 3.1 for further information). The current density (normJ) was used as recommended in the SimNIBS pipeline for our type of data (Saturnino et al., 2019b). In any case, it is worth to consider that normJ and normE distributions are equivalent, as J and E are directly connected via Ohm’s law, being the current density directly proportional to the electric field. Therefore, if not otherwise specified, figures and analysis results reflect the electric current density in all cases, and are expressed as amperes per square meter (A/m^2^). Due to technical issues during the generation of the head mesh, related to poor quality of the T1 MRI data, a subject was discarded from these analyses.

### 2.8 Statistical analyses

Data analyses were performed using IBM SPSS (IBM Corp. Released 2017. IBM SPSS Statistics for Windows, Version 25.0. Armonk, NY: IBM Corp.) and MATLAB (Version R2019a, The MathWorks Inc., Natick, MA, USA).

Rs-FC correlation matrices, permutation testing and pixel correction for multiple comparisons were performed using custom made MATLAB scripts. Functional connectivity differences were compared between conditions using non-parametric permutation testing (Nichols & Holmes 2002). We chose this method because it does not rely on assumptions about the distribution of the data and correction methods for multiple comparisons can be easily implemented (Theiler et al., 1992).

Time-series data for each subject, ROI and condition were concatenated into a 4D array (i.e., subject × time-point × ROI × condition). Then, correlations between all ROIs for each subject and condition were computed, taking time-points as individual observations in each ROI-to-ROI correlation. This resulted in three correlation matrices (i.e., one for each condition; Fig. 3A) with three dimensions each (i.e., ROI × ROI × subject). Then, differences between the means of the correlation matrices, for each pair of conditions, were computed (Fig. 3B) and compared with the null-hypotheses distribution generated by randomly shuffling the condition labels over subjects and repeating this procedure for 1000 iterations (Fig. 3C, supra-threshold: in yellow). The resulting comparison matrices were corrected for multiple comparisons using pixel correction (Cohen, 2014; Fig. 3C, pixel-corrected: in red). This procedure consists in picking the largest and smallest test statistic values of each permutation—which results in two distributions of extreme values—and then setting the thresholds for statistical significance to be the values corresponding to the 97.5 percentile of the largest value and the 2.5 percentile of the smallest value. Pixel correction was preferred as compared to the more common cluster-based methods because the nature of our data (i.e., correlation matrices) does not require that significant differences are spatially clustered.

**Fig. 3.**
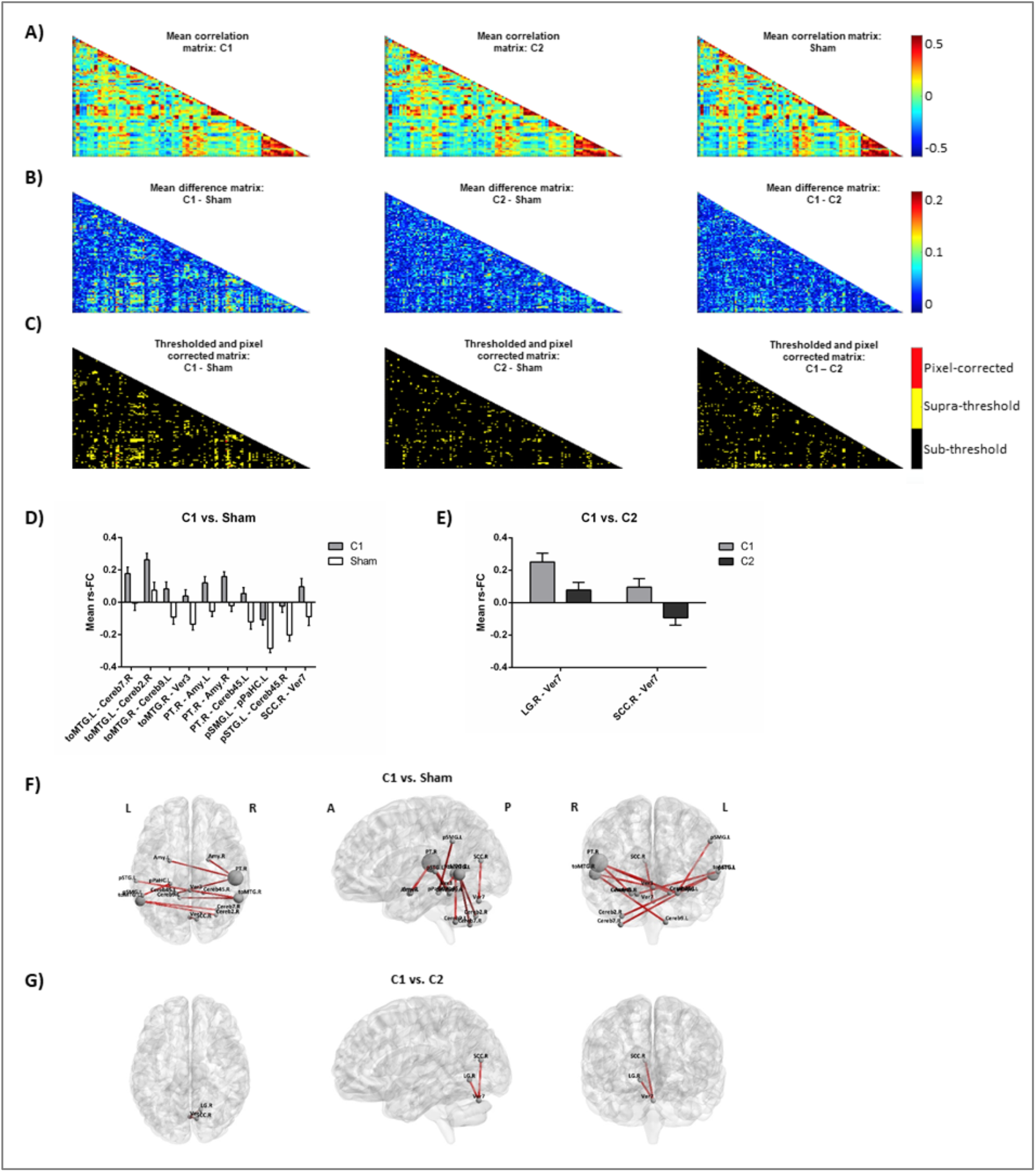
Rs-fMRI analyses. A) Mean correlation matrices for (left to right): C1, C2 and sham. B) Mean difference matrices for (left to right): C1 – Sham, C2 – Sham, C1 – C2. C) Thresholded and pixel corrected matrices for (left to right): C1 – Sham, C2 – Sham, C1 – C2 (dark: sub-threshold; yellow: supra-threshold; red: pixel-corrected). Bar-plots showing the mean of rs-FC comparing D) C1 vs. Sham and E) C1 vs. C2. Representation of the significant connections for F) C1 vs. Sham and G) C1 vs. C2 on a standard map (left to right: axial, sagittal and coronal view). Abbreviations: toMTG, Middle Temporal Gyrus, temporo-occipital part; Cereb, Cerebellum; Ver, Vermis; PT, Planum Temporale; Amy, Amygdala; pSMG, Supramarginal Gyrus, posterior division; pPaHC.L, Parahippocampal Gyrus, posterior division, pSTG, Superior Temporal Gyrus, posterior division; SCC, Supracalcarine Cortex; L, Left; R, Right; A, anterior; P, posterior.

Furthermore, overall electric field magnitude differences between C1 and C2 were compared using a paired sample t-test. Three additional paired sample t-tests were used to test for field magnitude differences between conditions for each of the three main ROIs selected based on rs-fMRI analyses. Finally, Pearson product-moment correlation was used to explore the association between NIBS-induced rs-fMRI effects and baseline rs-fMRI and modeling-based electric field magnitude estimates in the selected ROIs. No adjustment for multiple comparisons was applied in these exploratory statistical analyses. All tests were two-tailed and α was set at 0.05.

## 3. Results

### 3.1 Multifocal tDCS effects on rs-fMRI

The two tDCS conditions modulated connectivity at the rs-fMRI level differently. When comparing C1 against sham, numerous connections (i.e., 10 resting-state couplings) significantly increased their coactivation (Fig. 3D and 3F). A large number of the affected connections involved temporal and temporo-occipital areas and distinct cerebellar regions. In particular, three temporal areas emerged as main hubs (i.e., those with ≥ 2 significant rs-FC couplings). These regions were the toMTG.L and toMTG.R and the PT.R, which fall in the posterior part of the temporal lobe. When contrasting C1 against C2, two connections were detected to be significantly different (Fig. 3E and 3G). These results represented occipital-cerebellar couplings. Particularly, we observed significant modification of the connections between the right supracalcarine cortex (SCC.R) and the right lingual gyrus (LG.R) and the seventh lobule of the vermis (Ver7). No differences were observed between C2 and sham.

### 3.2 Estimated electric fields

The mean and *SDs* of electric fields induced by C1 and C2 are displayed in Fig. 4A (for C1) and 4B (for C2). Individually modeled electric fields are displayed in Fig. S3 (for C1) and S4 (for C2). As can be observed, the electric field induced by C1 predominantly affected the posterior part of the temporal lobule and its related temporo-occipital areas. In contrast, electric field induced by C2 showed a more anteriorly-centered distribution encompassing principally the precentral, superior and, in a lesser extent, the middle frontal gyri in a right-lateralized fashion. Moreover, as expected for the C2, the electric field magnitude within the posteromedial and occipital areas was very low. Thus, the intended original stimulation pattern and the obtained average in our subjects are fairly consistent (i.e., Fig. 2 and Fig. 4).

**Fig. 4.**
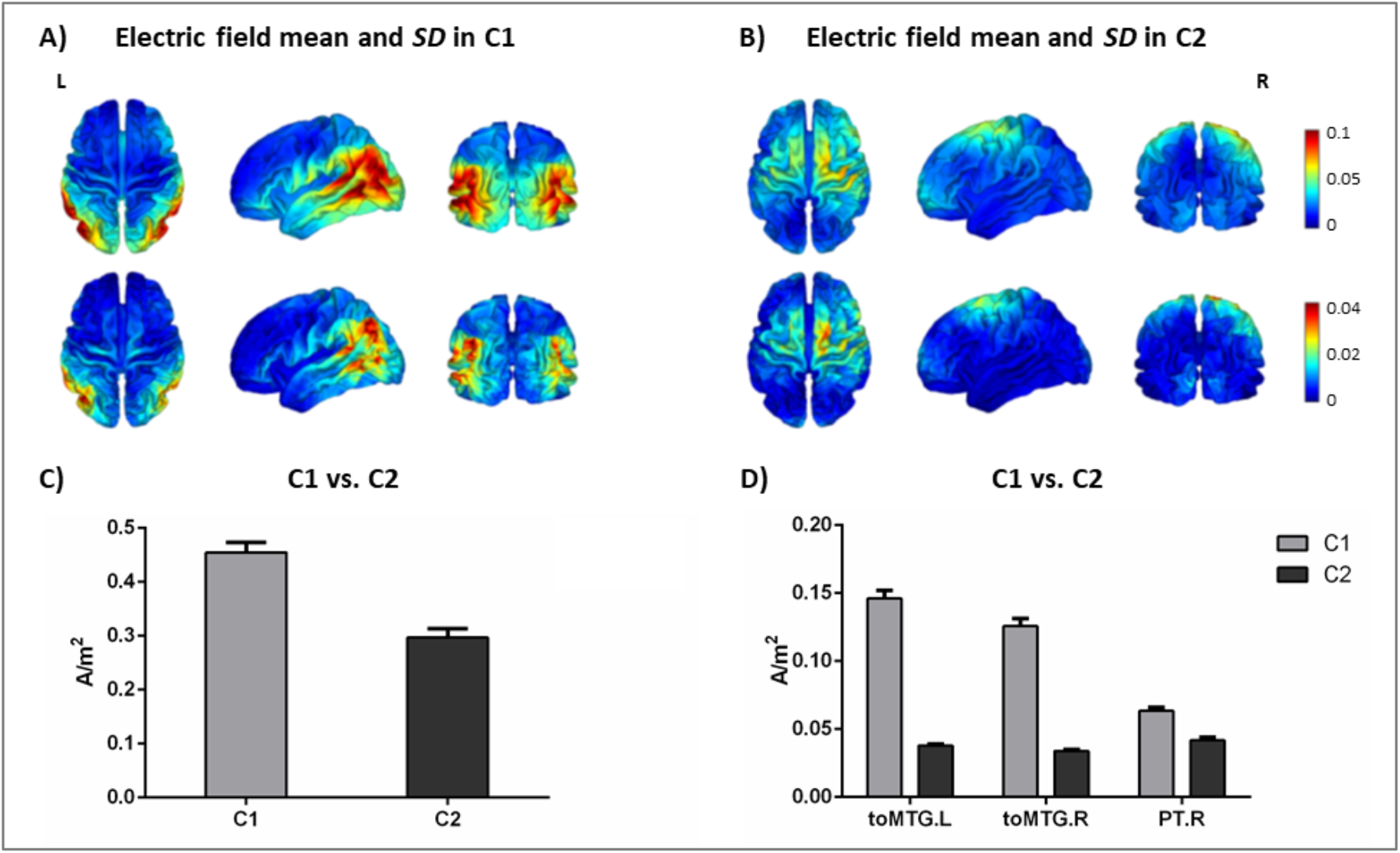
Multifocal tDCS estimated electric fields. Distribution of the average electric field distribution (top) and *SD* (bottom) for all subjects in A) C1 and B) C2 (in A/m^2^ units). C) Comparison between C1 and C2 of the mean’s magnitude of the top percentiles (99.9%) of the field (in A/m^2^ units). D) Comparison between C1 and C2 of the electric field magnitude extracted from the three main ROIs detected on the rs-fMRI analyses (i.e., toMTG.L, toMTG.R and PT.R; in A/m^2^ units). Abbreviations: C1, montage 1; C2, montage 2; toMTG, Middle Temporal Gyrus, temporo-occipital part; PT, Planum Temporale; L, Left; R, Right.

Overall, C1 induced a statistically higher field magnitude values than C2 when considering the mean of the top percentiles (99.9%) of the estimated field (*t* = 8.716; *p* < 0.001; Fig. 4C). Additionally, the electric field magnitude values extracted from the three main ROIs identified on the rs-fMRI analyses (i.e., toMTG.L, toMTG.R and PT.R), were, as expected, significantly higher in C1 when compared to C2 (toMTG.L: *t* = 19.986, *p* < 0.001; toMTG.R: *t* = 18.473, *p* < 0.001; PT.R: *t* = 10.275; *p* < 0.001; Fig. 4D).

### 3.3 Predictors of rs-fMRI modulation

Since no significant differences were observed between C2 and sham we restricted this part of the analysis to C1, which was significantly different from sham.

First, we conducted associations between tDCS-induced rs-fMRI changes and baseline connectivity. All statistically significant rs-FC modulations comparing C1 with sham were negatively associated with resting-state connectivity during sham (i.e., baseline rs-fMRI): toMTG.L-Cereb7.R: *r* = −0.666, *p* < 0.001; toMTG.L-Cereb2.R: *r* = −0.667, *p* < 0.001; toMTG.R-Cereb9.L: *r* = −0.791, *p* < 0.001; toMTG.R-Ver3: *r* = −0.788, *p* < 0.001; PT.R-Amy.L: *rs* = −0.627, *p* < 0.001; PT.R-Amy.R: *rs* = −0.625, *p* < 0.001; PT.R-Cereb45.L: *r* = −0.773, *p* < 0.001; pSMG.L-pPaHC.L: *r* = −0.677, *p* < 0.001; pSTG.L-Cereb45.R: *r* = −0.630, *p* < 0.001; SCC.R-Ver7: *r* = −0.517, *p* = 0.003 (Fig. 5 A-J).

**Fig. 5.**
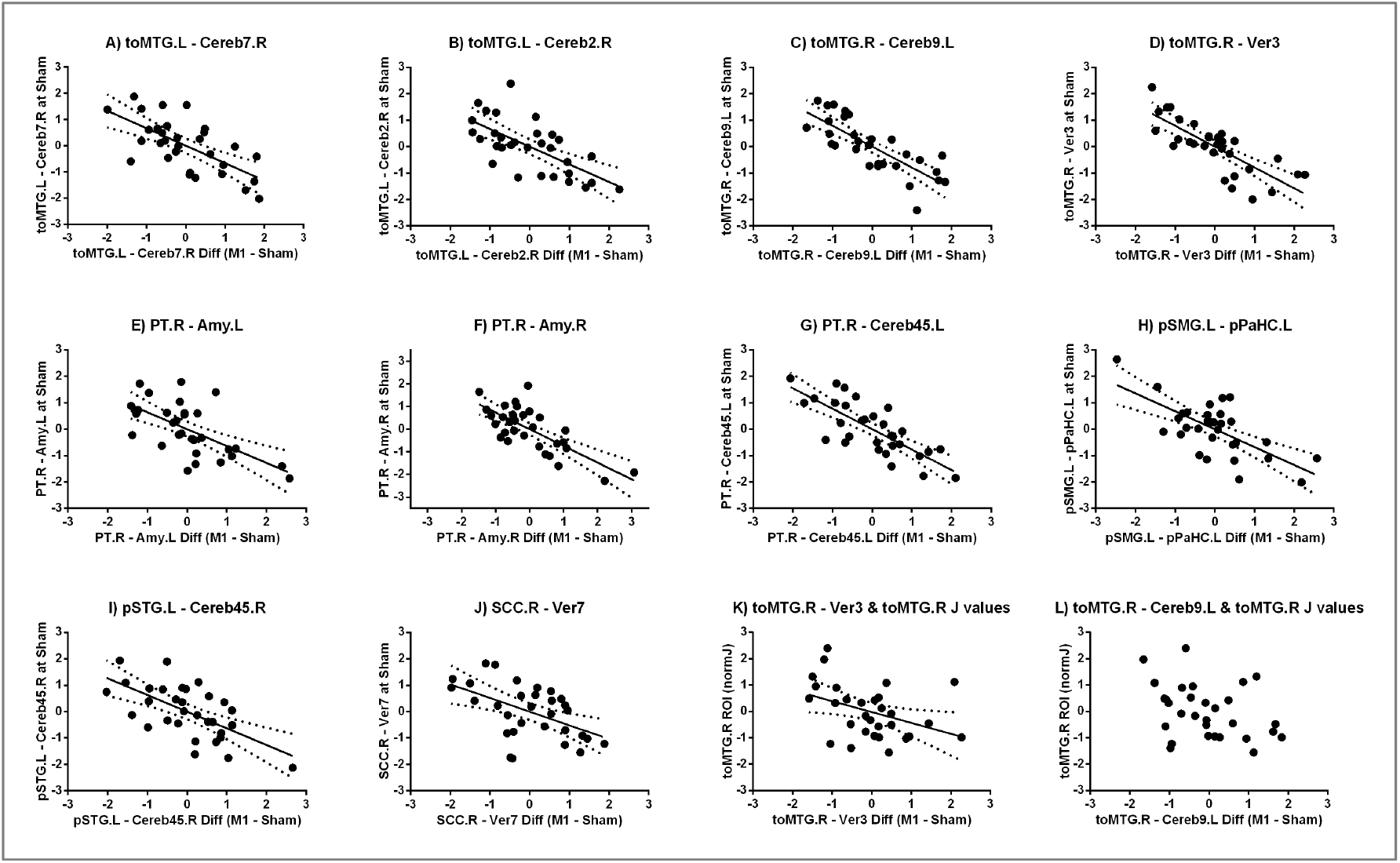
Predictors of tDCS-induced rs-fMRI responses. (A-J) Scatter plots showing the relationships between tDCS-induced changes in rs-fMRI and baseline rs-FC. (K-L) Scatter plots showing the relationships between tDCS-induced changes in rs-FC and electric field magnitude values. Data in A-L) are presented with *z* scores. Note that all the scatter plots are illustrated in the same scale (−3 to 3) but F), which goes from −3.5 to 3.5 for data distribution. Abbreviations: toMTG, Middle Temporal Gyrus, temporo-occipital part; Cereb, Cerebellum; Ver, Vermis; PT, Planum Temporale; Amy, Amygdala; pSMG, Supramarginal Gyrus, posterior division; pPaHC.L, Parahippocampal Gyrus, posterior division, pSTG, Superior Temporal Gyrus, posterior division; SCC, Supracalcarine Cortex; L, Left; R, Right.

Subsequently, correlation analyses between tDCS-induced rs-fMRI changes and calculated electric field magnitude estimates were performed. We observed that those subjects who showed increased coactivation in C1 compared to sham in the toMTG.R – Ver3 coupling also presented lower induced electric field magnitude values in C1 (*r* = −0.401, *p* = 0.028; Fig. 5K). In addition, those subjects who showed higher coactivation at C1 compared to sham in the toMTG.R – Cereb9.L also presented a negative association with the induced electric field magnitude estimates at C1, however, this was not statistically significant (*r* = −0.358, *p* = 0.052; Fig. 5L).

## 4. Discussion

This represents the first study investigating the impact of two distinct multifocal tDCS montages on rs-fMRI in healthy aging. Our results showed that: (1) multifocal tDCS modulates rs-FC in a montage-dependent manner in older adults. (2) Moreover, the functional impact is consistent with the spatial distribution of estimated electric field induced on the brain. (3) At the individual level, tDCS-induced rs-fMRI responses can be predicted by baseline connectivity and induced electric field magnitude estimates.

In this investigation, we observed that multifocal tDCS can modulate brain activity as measured through rs-fMRI level amongst older adults. Furthermore, network-based tDCS appears to modulate functional connectivity in a manner dependent on the estimated electric field. In this vein, the impact of transcranial stimulation was particularly evident when comparing C1 to sham. Three cortical areas emerged as the principal modulated regions: the left and right toMTG and the right PT. A relevant modulatory effect between these cortical nodes and the cerebellum was detected, though the modulation of cerebellar areas was less specific. In addition, in other cortical temporo-occipital areas, such as the posterior STG and SMG, and particular subcortical regions, such as the amygdala and hippocampal formation, rs-FC was also modulated. The main rs-fMRI results in C1 vs. sham (i.e., considering the main hubs; Fig. 3F) correspond anatomically with the estimated electric field induced on the brain, as the largest electric field simulated values in C1 were observed also in temporo-occipital brain regions (see Fig. 4A). Furthermore, these results were consistent with the position of the electrodes that delivered the highest stimulation intensities in C1 (Fig. 2E), namely P3 and P4 (with positive polarity), as well as P7 and P8 (with negative polarity). Interestingly, P3 and P4 electrode positions in the international 10-10 system are located above the angular Brodmann’s area 39 (BA39; Koessler et al., 2009), which is part of the parietal cortex, lying close to the junction of temporal, occipital, and parietal lobes. Furthermore, the P7 and P8 electrodes are situated above the temporo-occipital Brodmann’s area 37 (BA37; Koessler et al., 2009). Therefore, taking into account that the most powerful electrodes in the C1 montage (i.e., ~ 1.2 – 1.7 mA) likely drive the field distributions and that the areas most significantly modulated in our study match the ones under these electrodes, one could reasonably assume that in multifocal tDCS protocols, electrodes with very low current intensities (i.e., ~ 0.3 −0.7 mA) might be dispensable. Anyhow, our data show that they do not seem to induce any estimable or relevant effect at the rs-fMRI level in older populations. To sum up, regarding the C1 montage, we observed a clear linear relationship between the electrode’s assembly and injected current intensities, the corresponding simulated electric field distribution in the cortical surface, and the neuroimaging results obtained with rs-fMRI.

In previous literature, considerable individual variability in response to distinct NIBS protocols has been reported (Hamada et al., 2013; López-Alonso et al., 2014; Martin-Trias et al., 2018; Perellón-Alfonso et al., 2018; Wiethoff et al., 2014). These reports highlight the importance of identifying the individual predictors of NIBS effects, particularly in older adults, where substantial attempts have been made to modulate brain function and cognition. In the present study we focused on two potential factors contributing to such variability: (1) baseline rs-fMRI and (2) estimated electric field magnitude.

Regarding baseline rs-fMRI connectivity, we observed that for C1 it predicted, in a negative fashion, the tDCS-induced effects on rs-FC. Numerous studies have revealed that baseline functional connectivity regulate the response to NIBS procedures both in younger (i.e., Cárdenas-Morales et al., 2014; Nettekoven et al., 2015; Vidal-Piñeiro et al., 2015) and older samples (i.e., Abellaneda-Pérez et al., 2019b; Antonenko et al., 2019b). Our results are in line with some of this previous research claiming that those subjects with higher indices of coactivation at baseline tend to show lower NIBS-induced functional modulations. In this vein, Nettekoven et al. (2015) used a repetitive transcranial magnetic stimulation (TMS) protocol and classified participants on the basis of an increase of ≥ 10% in motor evoked potential amplitude, as either responders or non-responders. Next, authors observed that both groups presented different functional connectivity profiles, with responders having lower levels of baseline rs-fMRI when compared to non-responders. It has been argued that this data might reflect a ceiling effect underlying non-responsiveness to stimulation at the systems level (Nettekoven et al., 2015). Similarly, our study revealed that this association also exists in multifocal tDCS when applied in older adults.

Regarding the second potential NIBS predictor, we investigated whether dissimilar individual differences in rs-fMRI modulation were related to the electric field magnitude estimates within the designated main ROIs. We observed that the lower the rs-fMRI modulation, the higher the induced electric field strength. This is in agreement with recent results from Antonenko et al. (2019a), who reported the same inverse association in the motor cortex of younger adults. Also, in this line, studies using a variety of other NIBS protocols described similar associations between induced electric fields and detected NIBS effects. Likewise, Cabral-Calderin et al. (2016) reported that for particular transcranial alternating current stimulation conditions, electric field strength predicted functional connectivity changes. Moreover, using TMS, Mikkonen et al. (2018) observed a correlation between the estimated electric fields by tDCS with resting motor thresholds that were specific to the hand area of the primary motor cortex, showing stronger induced fields in individuals with lower motor thresholds. Our results further replicate these findings while providing first evidence for a relationship between estimated induced fields on the brain and the real neurophysiological effects in older adults. In addition, we have furthermore shown that induced electric field magnitude estimates have predictive value regarding NIBS effects in aging, most likely because these are contingent to the individual head and brain anatomy (Miranda et al., 2013; Thielscher et al., 2011).

Altogether, our data show that multifocal non-invasive stimulation models and protocols entail capability to effectively modulate precise fMRI configurations in older adults. These observations may have important future implications for the cognitive neuroscience of aging field, since many of the neural basis regarding cognitive functioning, longitudinal trajectories and inter-individual differences, including the development of theoretical models, have been mainly based on studies employing this imaging technology (Cabeza et al., 2018; Grady, 2012; Vaqué-Alcázar et al., 2020). Furthermore, fMRI changes can effectively track the positive impact of behavioral interventions aimed to ameliorate cognition in the elderly (Duda & Sweet, 2019), including its combined effects with NIBS (Antonenko et al., 2018). Therefore, and as opposed to the use of conventional tDCS montages, the possibility of *a priori* designing and modelling particular NIBS-based interventions incorporating individual information (i.e., baseline patterns of brain connectivity) that are predictive of regional-specific fMRI changes may offer a valuable and refined, individually-tailored, approach for studies intended to optimize brain functionality amongst older individuals, or to investigate the impact of interventions in clinical trials.

## 5. Limitations

The present study is not without limitations. First, it is worth noting that our original modellings designed with Stimweaver were based on the template Colin27. Models based on aged brains or on individual MRIs could have improved the electrodes montage and delivered stronger and more consistent effects for both montages. Another important limitation regarding the optimization is that it was conducted assuming that PiStim electrodes (Ag/AgCl 1 cm radius cylindrical electrodes with conductive gel underneath) were used in the montage. The electrodes that were actually employed were slightly larger (1.6 cm radius for the saline soaked sponge). Since electrode geometry has an influence on electric field distribution, it is likely that the optimized montage would have been different (both in terms of positions of the electrodes and their currents) had these electrodes been modeled during the optimization instead. Furthermore, although the optimization was focused on the normal component of the electric field, the magnitude was evaluated in the paper. This is because for our research purposes we consider net magnitude values more straightforward to interpret than those including current directionality data. In any case, future work should be conducted in determining the mechanisms for interaction between the electric field and the neurons, to determine the most reliable component(s) that underlie the effects of stimulation. Finally, it must be recognized that in all simulation-based electric field planning systems, there is uncertainty about the precise conductivity values that should be used for the different tissues and materials when creating and analyzing a model for a particular montage (see for further detail Miranda et al., 2013; Saturnino et al., 2019a). Notwithstanding, notable advances are being achieved in this field (Huang et al., 2017; 2018).

## 6. Conclusions

Present results highlight that multifocal electrical stimulation protocols are capable of modulating neural dynamics in the elderly. Moreover, this modulation is consistent with the anatomical distribution of the estimated electric fields on the brain. Thus, applying network-based procedures seems to be a novel feasible approach to accurately target and modulate specific fMRI networks and connections critically involved in the cognitive aging process. Finally, we have shown that specific factors partially accounting for the individual variability to NIBS are also relevant in aged populations. Gathering knowledge about these variables will allow us to ultimately refine the parameters of transcranial stimulation to boost the brain and cognitive benefits derived from NIBS-based interventions in advanced age.

## Supporting information

Abellaneda_Perez_et_al_2020_SM

## Acknowledgments

We are indebted to the Magnetic Resonance Imaging Core Facility of the IDIBAPS and the SimNIBS developers for the technical help.

## Data Availability Statement

The data that support the findings of this study are available from the corresponding author upon reasonable request.

