## Supplementary material for "Multifocal tDCS modulates resting-state functional connectivity in older adults depending on induced electric field and baseline connectivity": Abellaneda_Perez_et_al_2020_SM

### Materials and methods

#### Neuropsychological assessment

A comprehensive battery of neuropsychological tests covering all cognitive domains was administered, including the Rey-Osterrieth Complex Figure (ROCF), Rey Auditory Verbal Learning Test (RAVLT), Boston Naming Test (BNT), Semantic category evocation of animals, Number location and incomplete letters from the Visual Object and Space Perception Battery (VOSP), Trail Making Test (TMT), parts A and B, Phonemic fluency (FAS), Stroop Color Word Test, Symbol Digit Modalities Test (SDMT), and Digit span forward and backward from WAIS-III. Finally, the Vocabulary Subtest from WAIS-III was also administered to have a measure of premorbid intelligence.

#### tDCS parameters

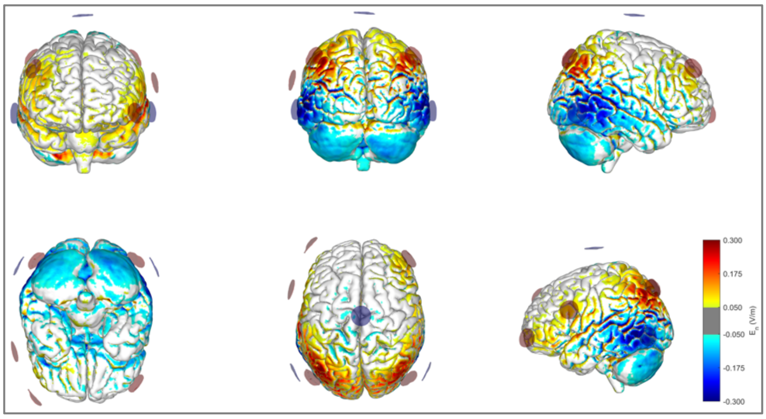


**Fig. S1.** Distribution of the $E_n$ field in the cortical surface in C1 (in V/m units).

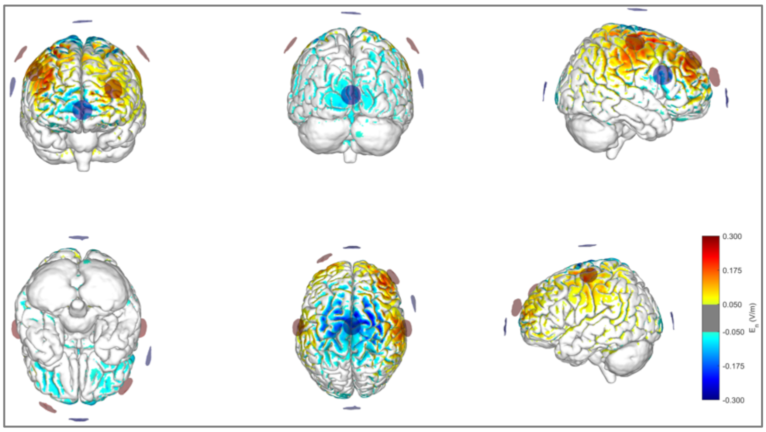


**Fig. S2.** Distribution of the $E_n$ field in the cortical surface in C2 (in V/m units).

### **Results**

#### Estimated electric field distributions

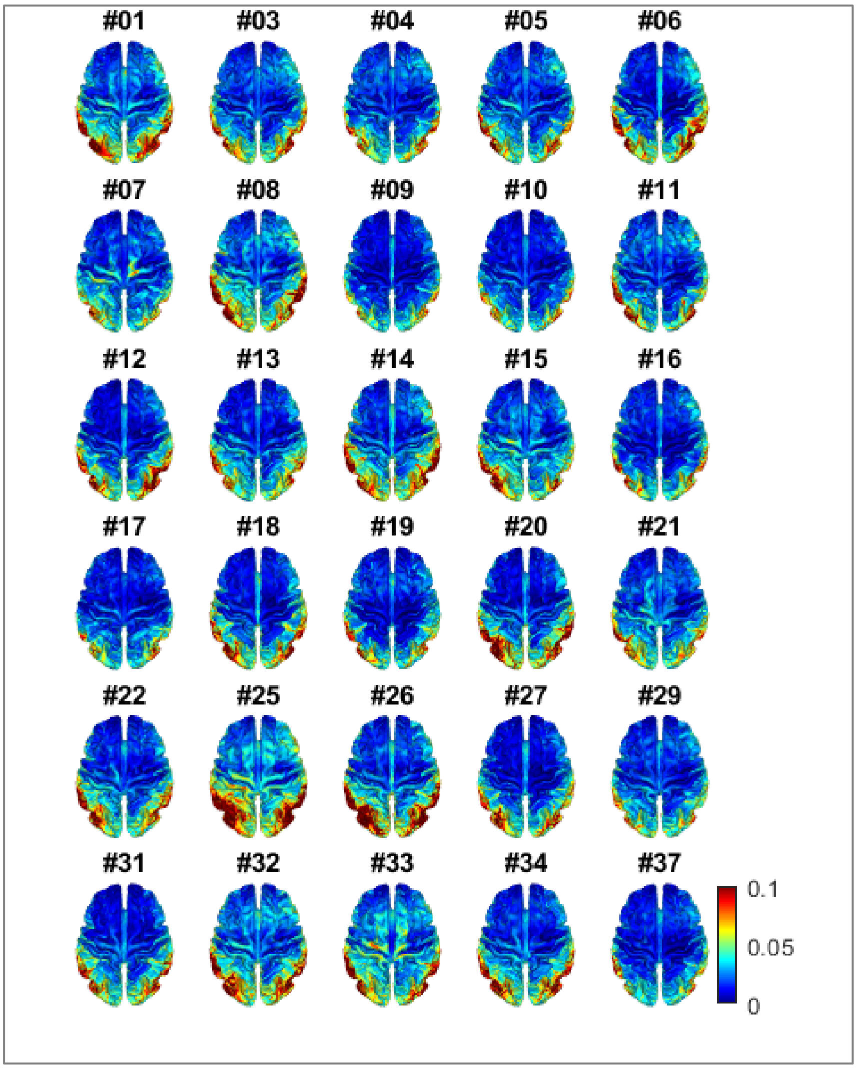


**Fig. S3.** Axial view of the norm component of the electric field simulations computed in each subject in C1 (in A/m$^2$ units). The # represents the identification of included subjects in these analyses in correlative order.

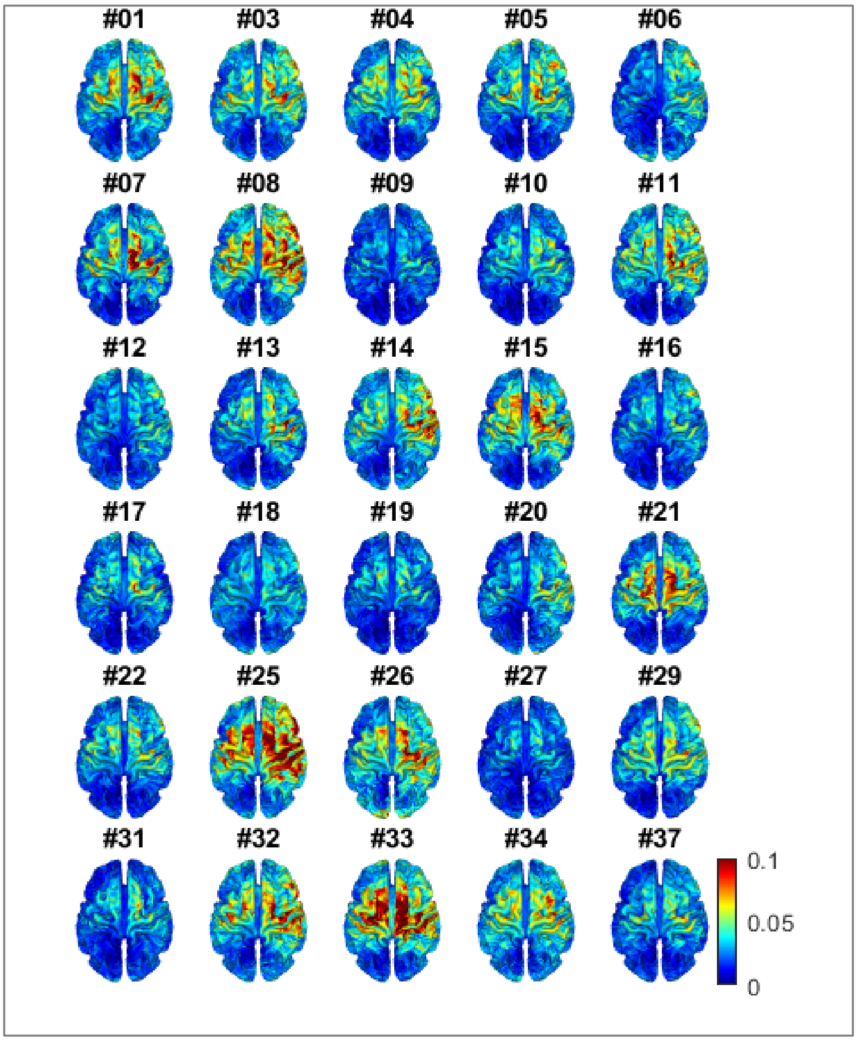


**Fig. S4.** Axial view of the norm component of the electric field simulations computed in each subject in C2 (in $A/m^2$ units). The # represents the identification of included subjects in these analyses in correlative order.
